# Architecture and assembly dynamics of the essential mitochondrial TIM chaperone systems

**DOI:** 10.1101/2020.03.13.990150

**Authors:** Katharina Weinhäupl, Yong Wang, Audrey Hessel, Martha Brennich, Kresten Lindorff-Larsen, Paul Schanda

## Abstract

The mitochondrial Tim chaperones are responsible for the transport of membrane proteins across the inter-membrane space to the inner and outer mitochondrial membranes. TIM9·10, a hexameric 70 kDa protein complex formed by 3 copies of Tim9 and Tim10, guides its clients across the aqueous compartment. The TIM9·10·12 complex is the anchor point at the inner-membrane insertase complex TIM22. The subunit composition of the TIM9·10·12 complex remains debated. Joint NMR, small-angle X-ray scattering and MD simulation data allow us to derive a structural model of the TIM9·10·12 assembly, which has a 2:3:1 stoichiometry (Tim9:Tim10:Tim12). We find that both TIM9·10 and TIM9·10·12 hexamers are in a dynamic equilibrium with their constituent subunits, exchanging on a minutes time scale. Residue-resolved NMR data establish that the subunits exhibit large conformational dynamics: when the conserved cysteines of the CX_3_C-X_*n*_-CX_3_C motifs are formed, short marginally stable *α*-helices are formed, and these are fully stabilized only upon formation of the mature hexameric chaperone. We propose that the continuous subunit exchange is a means of mitochondria to control their level of inter-membrane space chaperones, and thus rapidly adapt to the cellular state.

## Introduction

Mitochondria perform a myriad of functions and are involved in most major metabolic pathways in eukaryotes. Of the more than 1000 mitochondrial proteins, ca. 99% are synthesized by cytosolic ribosomes and imported as precursor proteins by dedicated chaperones, receptors, translocases and insertases (1–3). This orchestrated transport and insertion process is particularly demanding for precursors of membrane proteins because their hydrophobicity makes them highly aggregation-prone. The family of small Tim chaperones ensures the transport of membrane precursor proteins from the central entry pore in the outer membrane, TOM40, across the inter-membrane space to the insertases located in the inner and outer membranes (4–7). The small Tim family in yeast comprises five proteins, Tim8, Tim9, Tim10, Tim12 and Tim13, each of ca. 10 kDa molecular weight, of which Tim9, Tim10 and Tim12 are essential. Strictly conserved CX_3_C-X_*n*_-CX_3_C motifs in all small Tims form two disulfide bonds that bridge the two CX_3_C sequences. Three hexameric TIM complexes have been identified *in vivo*, TIM9·10, TIM8·13 and TIM9·10·12. Their interaction with membrane precursor proteins has been studied in detail by *in vivo* and biochemistry approaches (8–10), and its atomic-resolution mechanism has been resolved recently (11).

High-resolution crystallographic structures of TIM9·10 and TIM8·13 (12–14) revealed an alternating arrangement of three copies of each subunit type (Tim9/Tim10 or Tim8/Tim13). Each subunit comprises an N-terminal and a C-terminal *α*-helical tentacle with flexible ends. Formation of the two intra-molecular disulfide bonds in each subunit stabilizes the helix-turn-helix fold of the assembled chaperones.

TIM9·10·12 is part of the TIM22 insertase in the inner membrane. No crystal structure of the TIM9·10·12 complex (Tim9:Tim10:Tim10b in humans) has been solved, and the number and arrangement of the three different types of subunits had remained elusive for long time. Based on sequence homology (15) or observation of interaction of Tim12 and Tim9 monomers (16) and import assays (17), it has been proposed that Tim12 replaces one or several copies of Tim10 to form the TIM9·10·12 complex. Only very recently two cryo-EM structures of the TIM22 from yeast (18) and human (19) have revealed the structures of the TIM9·10·12 complex at 3.7 to 3.8 Åresolution. Intriguingly, the structures modeled into the two cryo-EM maps reveal different stoichiometries with Tim9:10:12 ratios of either 2:3:1 (human) or a 3:2:1 (yeast). This apparent variability between different species is remarkable, given that the orientation and arrangement of the hexameric chaperone within the two TIM22 complexes were essentially identical. This observation raises the question about inherent flexibility of the stoichiometry, and about the stability of the assembly.

Here we use an integrated structural biology approach to provide detailed insight into the structure of the TIM9·10·12 complex, as well as the assembly process, and the subunit dynamics at the atomic level. We derived a structural model of TIM9·10·12, in which we show to have a 2:3:1 (Tim9:10:12) stoichiometry. We find that the hexameric chaperones, TIM9·10·12 as well as TIM9·10, are in continuous exchange with its constituent subunits, whereby the disassembled subunits are in equilibrium with hexamers. We show that the structure of the subunits depends both on the oxidation state of the disulfides and assembly state: a highly dynamic fully unfolded state exists when the disulfide bonds are not formed, and disulfide formation induces short, marginally stable *α*-helices. Only assembly to hexameric complexes stabilizes these *α*-helices fully, leading to a rigid core of the chaperone with flexible N- and C-terminal tentacles.

Our observation of continuous subunit exchange points to a mechanism by which the chaperones can be assembled from a pool of subunits, based on the mitochondrial import demand. We demonstrate that such subunit exchange persists even when physiological full-length client proteins are attached, highlighting the inherently dynamic character of these chaperone complexes.

## Results

### Isolated small Tim subunits exhibit large oxidation-dependent flexibility

*In vivo*, small TIMs are imported into mitochondria with their cysteines in the reduced state, and are then oxidized with the aid of Mia40 (20), and they finally assemble to the mature hexameric chaperones. To understand the structural consequences of disulfide formation, we investigated the structure and dynamics of Tim9, Tim10 and Tim12 in the reduced and oxidized states using solution-state nuclear magnetic resonance (NMR) spectroscopy, which is exquisitely sensitive to local structure and flexibility at a resolution of individual atoms. Early reports of NMR spectra indicated a partial molten globule character of Tim9 and Tim10 (21), but they lacked site-specific assignments, making conclusions difficult.

We prepared separate samples containing only either Tim9, Tim10 or Tim12 by bacterial over-expression and purification, and performed NMR experiments in the presence and absence of the reducing agent tris(2-carboxyethyl)phosphine (TCEP), i.e., with the disulfide bonds broken or intact, respectively. We assigned Tim9, Tim10 and Tim12 in all available states, oxidized and reduced, as well as the hexameric state of TIM9·10 (11) and TIM9·10·12 (see below). The atom-wise NMR chemical shifts provide direct information about the local backbone dihedral angle, and thus about the secondary structure, in a residue-wise manner. Another parameter, the heteronuclear ^1^H-^15^N nuclear Overhauser enhancement (hetNOE), provides quantitative information of the amplitude of backbone motion on sub-nanosecond time scales, allowing to further identify the disordered regions. HetNOE values in folded rigid proteins are expected on the order of 0.6-0.8, while lower values indicate large-amplitude fast motions (picoseconds).

In the isolated subunits with reduced cysteines, all small Tims show NMR parameters characteristic of fully disordered polypeptide chains (Fig. 1). The α-helix propensity, derived from the assigned chemical shifts, is below 10% for the majority of residues, and a nuclear spin relaxation parameter, the ^1^H-^15^N heteronuclear nuclear Overhauser enhancement (hetNOE) values are around or below zero, indicative of large-amplitude sub-nanosecond motions (Fig. 1J and S1H,I).

**Fig. 1.**
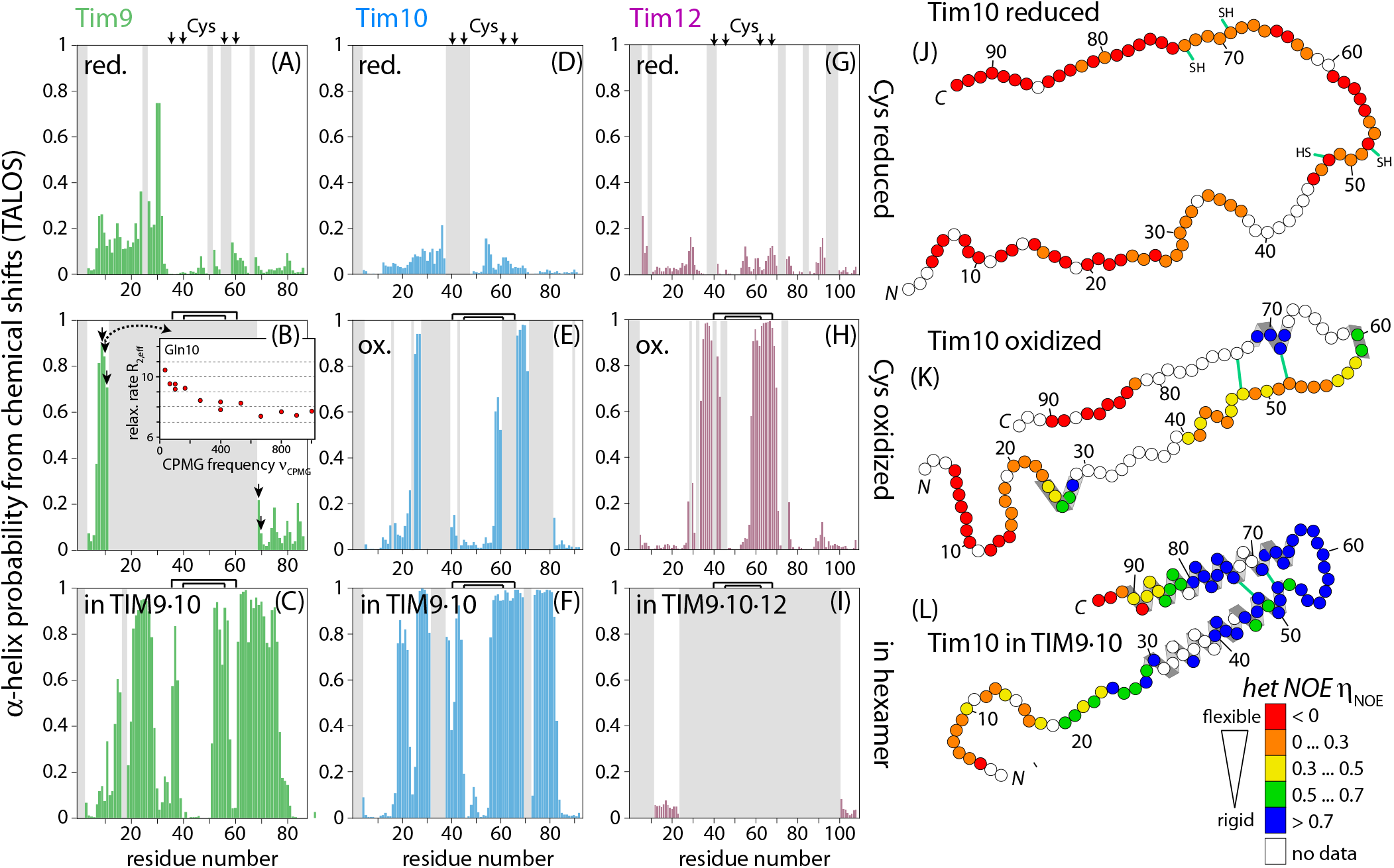
Isolated Tim subunits are highly dynamic, form short flexible *α*-helices only when the disulfides are formed, and fold only upon hexamer formation. (A)-(I) Residue-wise chemical-shift derived *α*-helix propensities for Tim9 (green), Tim10 (blue) and Tim12 (purple) in the three different sample states: isolated Cys-reduced (with TCEP), isolated oxidized (without TCEP) and oxidized hexameric. (J)-(L) Illustration of the ^1^H-^15^N heteronuclear nuclear Overhauser enhancement (hetNOE) values of the three states of Tim10, illustrating increasing rigidification from the reduced to the oxidized, asssembled state. Schematic helices are shown for the parts with a TALOS-derived α-helicity above 0.5, or which are in helix conformation in the crystal structure. The hetNOE data are plotted in Fig. S1H,I.

When the disulfide bonds are formed, the small Tims behave significantly different. A first striking observation is that the NMR signals of many residues become broadened beyond detection. The majority of the non-detected residues are located in regions which form *α*-helices within the assembled hexameric complex. A common reason for such NMR line broadening is the presence of conformational dynamics on the millisecond (ms) timescale, which we probed by CarrPurcell-Meiboom-Gill (CPMG) relaxation-dispersion (RD) NMR experiments (22). Non-flat RD profiles of the visible residues adjacent to the invisible residues (Fig. 1B, insert) directly reveal the presence of ms motions. Thus, we conclude that like in many molten-globule states (23), millisecond dynamics is present in part of the oxidized unassembled Tims. The chemical shifts of the observed residues in the oxidized unassembled Tims demonstrate the presence of *α*-helical backbone conformation in regions which form *α*-helices also in the hexameric TIM9·10 assembly. However, the secondary structure elements in the isolated Tim9, Tim10 or Tim12 are shorter than the ones in the hexameric state (Fig. 1). The presence of secondary structure in the isolated small Tims with intact disulfide bonds is also evident from the increased heteronuclear NOE values in the regions exhibiting *α*-helical backbone conformation. This observation directly shows that the monomers are less dynamic than in the reduced state.

Taken together, NMR analyses show that in the absence of disulfide bonds, Tim9, Tim10 and Tim12 behave as fully flexible random coils, while disulfide bonds induce short *α*-helical structures that undergo extensive millisecond motions reminiscent of molten globule states.

### Structure of TIM9·10·12 reveals 2:3:1 stoichiometry and flexible tails

We next interrogated the structure and dynamics of the hexameric TIM9·10·12 chaperone in solution. To gain insight into the TIM9·10·12 complex structure, we produced homogeneous and pure TIM9·10·12 complex from TIM9·10 and Tim12. Briefly, His-tagged Tim12 was immobilized on an affinity resin, and TIM9·10 was added, allowing to form TIM9·10·12 complex; excess TIM9·10 was eliminated before elution. Analytical ultracentrifugation, size-exclusion chromatography and NMR-detected translational diffusion experiments show that TIM9·10·12 forms a hexamer of similar size as TIM9·10, as expected (Fig. S3). The obtained TIM9·10·12 complex appeared to be independent of the relative amounts of TIM9·10 and Tim12 used in this pull-down procedure, and the stoichiometry did not further evolve when Tim12 was added to TIM9·10·12 after elution (data not shown), suggesting that under the experimental conditions this is a stable species.

A central question related to the TIM9·10·12 structure is its Tim9:Tim10:Tim12 stoichiometry and the spatial arrangement of subunits. We used NMR spectroscopy to unambiguously resolve this question. Central to this NMR approach is the realization that while TIM9·10 has a three-fold symmetry with alternating Tim9 and Tim10 subunits, the symmetry is broken in TIM9·10·12, because a given subunit type can have up to three different environments (Fig. 2A). Accordingly, in TIM9·10 we observe a single set of NMR resonances for the three symmetry-related Tim9 and Tim10 subunits, as each subunit has the same neighbors (11), while more peaks are expected in TIM9·10·12 complexes. Therefore, determining how many copies of a given subunit are present in the complex boils down to counting the number of peaks of a given residue in a given subunit type. The apparent peak multiplicity may be smaller than the number of non-equivalent subunits because of peak overlap, but the number of non-equivalent subunits dictates the maximum number of observable peaks.

**Fig. 2.**
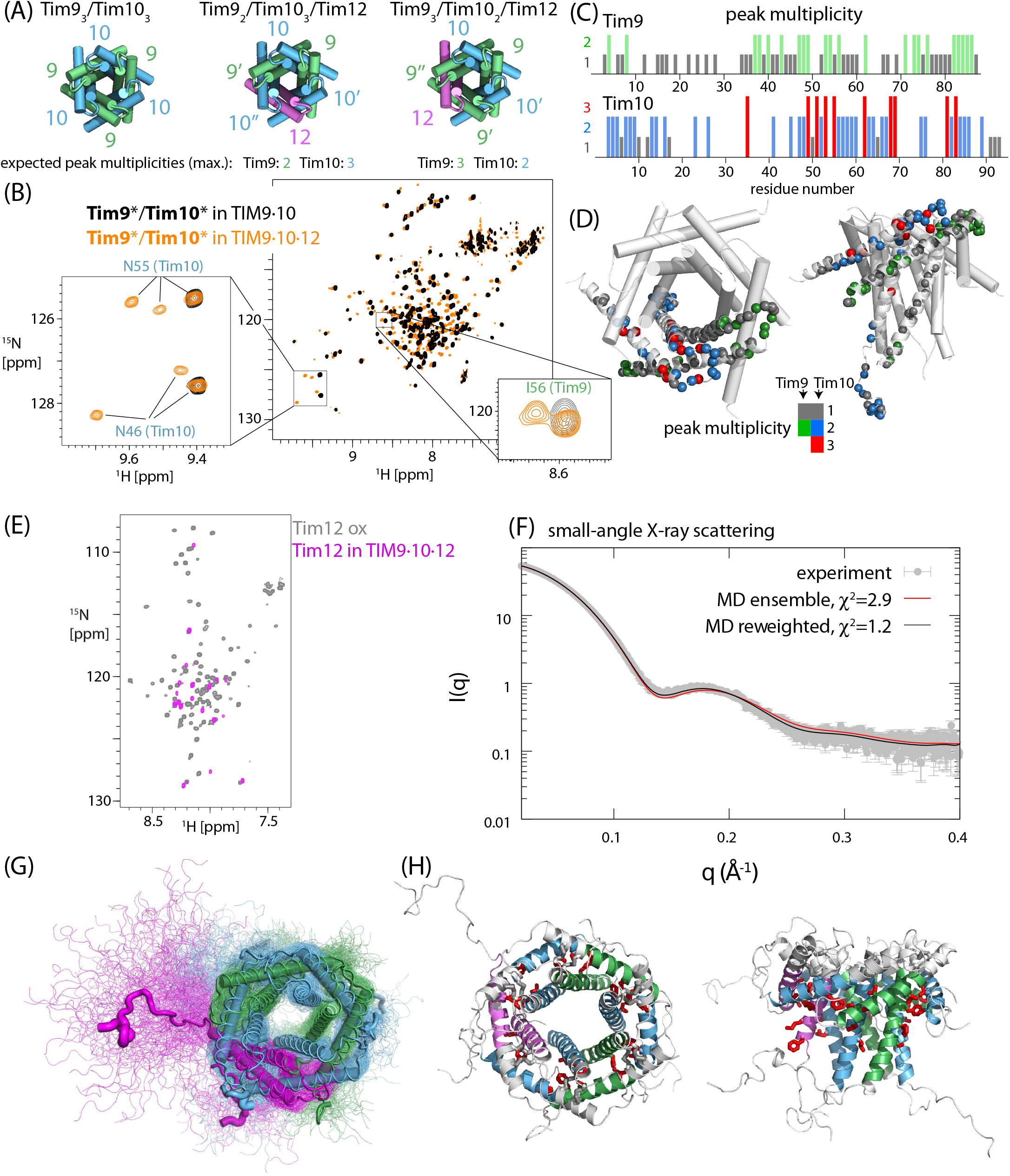
Structural model of TIM9·10·12 from NMR, SAXS and MD. (A) Symmetries of TIM9·10 and two possibilities of TIM9·10·12 with different stoichiometry, resulting in different numbers of peak multiplicities. (B) ^1^H-^15^N spectra of TIM9·10 and TIM9·10·12 samples, in which Tim9 and Tim10 subunits are ^2^H-^15^N-labeled. Two examples of tripled peaks in Tim10 are highlighted. (C, D) Number of observed ^1^ H-^15^ N peaks per residue. (E) ^1^ H-^15^ N spectra of ^2^ H-^15^ N-labeled Tim12 in isolation or in the TIM9·10·12 complex. (F) SAXS data of TIM9·10·12 and the back-calculated SAXS curve from the MD ensemble. (G) Conformational ensemble of TIM9·10·12 from MD which results in the black back-calculated curve of panel (F). Residue-wise helix propensities and RMS fluctuation from the MD simulation are shown in Fig. S4. (H) The conserved hydrophobic residues in TIM9·10·12, highlighted as red sticks on the SAXS/NMR/MD-derived structural model, point into the cleft between inner and outer tentacles.

We have prepared NMR samples of TIM9·10·12 in which either both Tim9 and Tim10 are ^2^H,^15^N-labeled and thus NMR visible (Fig. 2B), or in which only either Tim9 or Tim10 or Tim12 was isotope labeled (Figs. S2 and 2E, respectively). ^2^H-^15^N correlation spectra of these samples only report peaks from the labeled subunit(s), which facilitates counting of peaks, and assigning them to individual residues in either Tim9 or Tim10. An additional 3D HNCO experiment corroborated the assignment (Fig. S2). We ruled out the possibility that these additional peaks in TIM9·10·12 compared to TIM9·10 arise from the presence of disassembled subunits in solution, by comparison to the spectra of disassembled subunits (Fig. S2 C).

The obtained peak multiplicity per residue, shown in Figure 2C, reveals that the NMR signals of at least ten residues in Tim10 are tripled (fully spectrally resolved) and for 32 residues we detect two resolved peaks. In Tim9, 22 residues have spectrally resolved doubled peaks, but no residue shows three peaks. These findings unambiguously establish that there are three non-equivalent Tim10 subunits and only two of Tim9. The residues with tripled NMR resonances in Tim10, and with doubled signals in Tim9, are primarily located in the regions in direct contact with neighboring subunits, where the neighbor-induced differences in chemical shifts are most pronounced (Fig. 2C). Collectively, the observed peak multiplicities in Tim9 (up to 2) and Tim10 (up to 3) establish the 2:3:1 (Tim9:Tim10:Tim12) stoichiometry.

We also performed NMR analyses of ^2^H,^15^N-labeled Tim12 within TIM9·10·12. While Tim12 has significant homology with Tim9 and Tim10 for the first ca. 80 residues (sequence similarity of ca. 55 and 70%, respectively), Tim12 bears a C-terminal extension. As it has been proposed that this part may interact with membranes (24), the TIM22 insertase or other small Tims, it is of interest to characterize the structure of this C-terminal extension. The NMR signals of ^2^H,^15^N-labeled Tim12 within TIM9·10·12 are of low intensity, and only a few highly flexible parts are visible (Figs. 2E, 1). Low signal intensity is common for a 70 kDa complex; however, the fact that resonances of Tim9 and Tim10 in this complex are readily detected points to an additional mechanism of line broadening in Tim12, possibly motions on millisecond time scales. The observable parts of Tim12 comprise the interesting C-terminal part as well as a stretch towards the N-terminus. Both regions are in a dynamic random-coil conformation according to the chemical shifts (Fig. 1I), unambiguously ruling out the possibility that the C-terminal extension forms a highly-populated *α* -helical conformation in solution. We propose that peak broadening in the Tim12 subunit in TIM9·10·12 arises due to µs-ms mobility within the complex. In line with this observation, we note that in most cases of tripled Tim10 peaks, one peak has significantly lower intensity, and these peaks may correspond to the subunit adjacent to Tim12.

We turned to small-angle X-ray scattering (SAXS) to obtain insight into the structural envelope of the ensemble of conformers (Fig. 2F), and used molecular dynamics (MD) simulations to link all experimental observables to structural views of the complex. For the MD simulations, we built a model of TIM9·10·12 based on the crystal structure of TIM9·10 and the NMR-derived information about stoichiometry and subunit arrangement, and performed an all-atom 3 µs-long MD simulation. The SAXS curve computed from the ensemble of conformations sampled along the trajectory is shown in Fig. 2F. The SAXS curve from the MD ensemble is in excellent agreement with the experimental data. The SAXS data of TIM9·10·12 are detectably different from those of TIM9·10, with more long distances, which we ascribe to the extended tail of Tim12 (Fig. S3F). A view of the ensemble of conformations of TIM9·10·12 is shown in Fig. 2G and Supplementary Movie 1. The content of (*α*-,3_10_-and *π*-)helical conformation for all chains, averaged over the MD trajectory, is shown in Fig. S4. Interestingly, along the MD trajectory, the C-terminal extension of Tim12 remained in a highly flexible state, rather than forming a helical conformation, in agreement with the observed NMR chemical shifts. Interestingly, in the cryo-EM model of TIM22, a weak but detectable density was modeled as part of Tim12’s C-terminus, but the last 11 residues were absent (19).

Together, NMR, SAXS and MD allowed us to derive a realistic structural model of the hexameric TIM9·10·12 complex in solution with a 2:3:1 (Tim9:Tim10:Tim12) stoichiometry and flexible tentacles.

### The hexameric chaperone assemblies are in dynamic equilibrium with the free subunits

Intriguingly, in addition to the NMR signals corresponding to the hexameric (Tim9)_3_·(Tim10)_3_ assembly, the NMR spectra of TIM9·10 samples feature a second set of cross-peaks with an intensity of a few percent relative to the major hexamer peaks (Figs. 3A and S5). We could directly assign the additional peaks in the TIM9·10 sample to the C-terminal residues of both Tim9 and Tim10 (9 and 5 C-terminal residues assigned, respectively) in triple-resonance backbone assignment experiments. The positions of these peaks correspond to those that we had found in samples of isolated Tim9 or Tim10 subunits. We see primarily the C-terminal residues of the isolated subunits, in our hexamer samples, as those are the most intense peaks, and the signals of other parts of the monomeric species is most likely below detection threshold in NMR assignment experiments. Similarly to TIM9·10, we also find minor peaks in the TIM9·10·12 sample (Fig. S2 D).

**Fig. 3.**
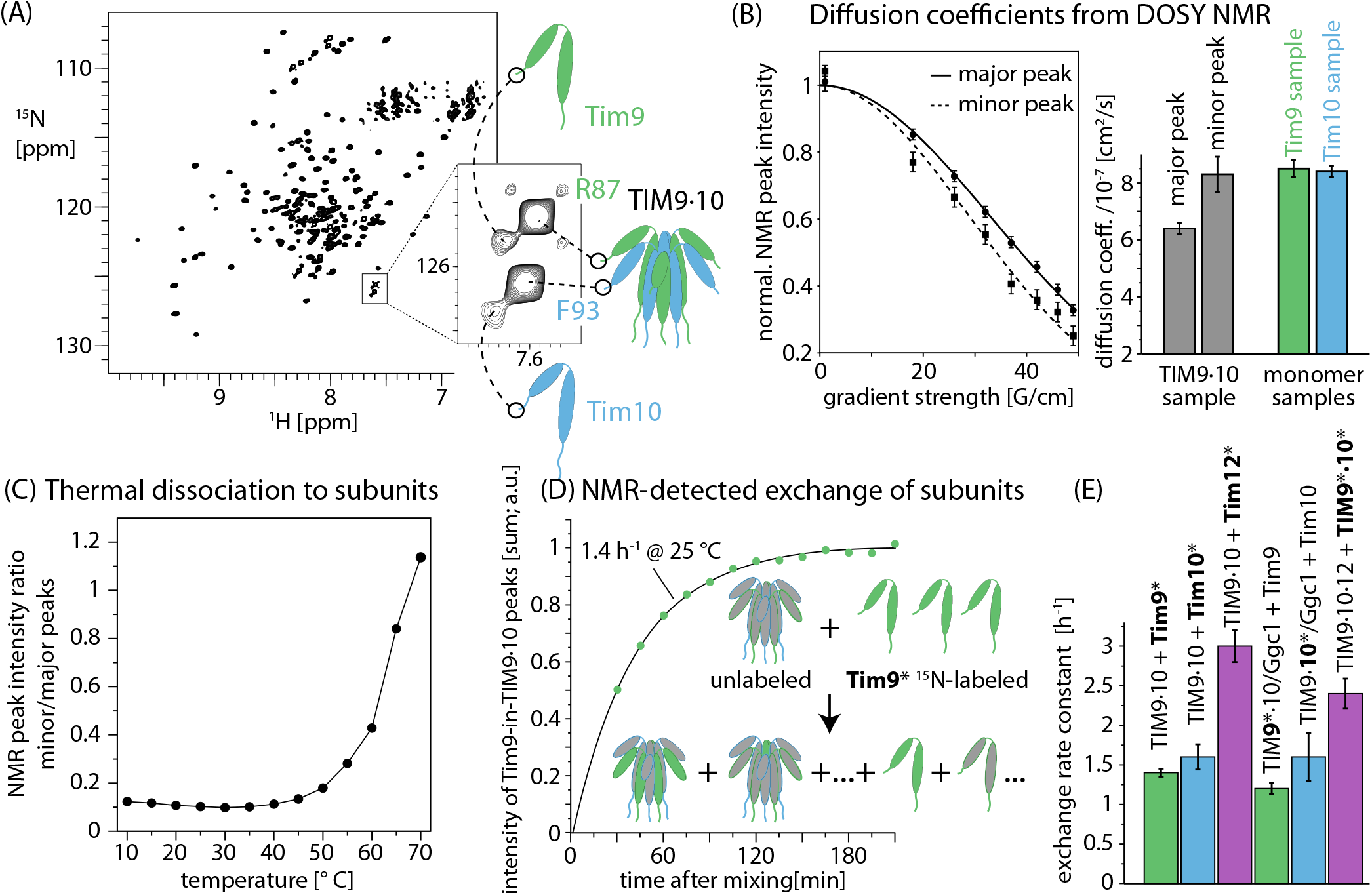
Tim subunits are in dynamic equilibrium between assembled and disassembled states. (A) ^1^H-^15^N NMR spectrum of TIM9·10, and a zoom onto the C-terminal residues, showing peak doubling. (B) Diffusion-ordered NMR curves of the sum of minor peaks and major peaks, and fitted diffusion coefficients. (C) Intensity ratio of minor over major peaks in TIM9·10 spectra as a function of temperature. (D) Kinetics of the buildup of Tim9 NMR ^1^H-^15^N peaks at the position of the TIM9·10 hexamer, after mixing unlabeled TIM9·10 with ^15^N-labeled Tim9. (E) Fitted kinetic rates of subunit exchange in different samples.

To get a direct measure of the size of these additional species, we measured the translational diffusion coefficients of these minor peaks by using a series of two-dimensional ^1^H-^15^N-edited diffusion-ordered NMR (DOSY) spectra on the TIM9·10 sample. The diffusion coefficients observed for the minor peaks match those measured in samples containing only Tim9 or Tim10, showing that the minor species in the TIM9·10 sample corresponds to free subunits (Fig. 3B). Temperature-dependent NMR spectra of TIM9·10 show that the peak heights of these C-terminal residues in the free subunits are about ten-fold lower than those of the TIM9·10 hexamer at ambient temperature, and that their intensity increases with temperature (Fig. 3C). Remarkably, even above 60*°*C, the hexamer signals are clearly visible. This process of population-shift to the unassembled species is fully reversible when decreasing the temperature: NMR spectra of hexameric TIM9·10 obtained after heat shock to 80 *°*C are indistinguishable from those of freshly co-purified TIM9·10. This reversibility points an equilibrium in which subunits of TIM9·10 can dynamically exchange with monomeric/dimeric species.

We investigated at which rate this dynamic subunit exchange occurs, and followed the incorporation of subunits into TIM9·10 by kinetic NMR experiments. Briefly, we prepared hexameric TIM9·10 complex by co-expression, as well as subunits of either Tim9 or Tim10, ensuring that these two samples (hexamer and isolated species, respectively) were differently isotope labeled (e.g., ^15^N-labeled Tim9 and unlabeled, i.e., NMR-invisible TIM9·10 hexamer). Upon mixing of these samples, the initial NMR spectrum is expected to contain only signals from labeled species, which is in its monomeric form at the onset of the reaction, and distributes over hexameric and monomeric species over time. Following the buildup of peaks corresponding to the hexamer and decay of monomer thus allows quantifying the reaction rate. Fig. 3D,E shows that subunits (Tim9, Tim10, Tim12) exchange on a time scale of tens of minutes.

We performed subunit exchange kinetic experiments also with chaperone complexes engaged with a full-length membrane protein client, the 30 kDa GDP/GTP carrier (Ggc1) (11). Interestingly, subunits are still able to enter/exit the complex when Ggc1 is wrapped around the chaperone (Figure 3E), albeit at a slightly reduced exchange rate constant. This finding underlines the highly dynamic nature of the chaperone-precursor protein complex.

Taken together, we find that Tim9, Tim10 and Tim12 can spontaneously insert into hexameric complexes and replace subunits from the hexamer.

## Discussion

We have used an integrated approach including NMR, SAXS, biophysics experiments and molecular dynamics simulations to understand the properties of the chaperone subunits in different states, to resolve the structure of the TIM9·10·12, and to bring to light the subunit exchange.

The TIM9·10·12 structure we have reported unambiguously establishes the stoichiometry of 2:3:1 (Tim9:10:12) in solution. This stoichiometry agrees with the one observed in the human TIM22 complex (19), but differs from the one fitted into the yeast TIM22 EM map (18). At a resolution of 3.8 Å, determining the identity of side chains unambiguously is generally not possible, and model building is, thus, challenging; however, support for the modeled structures comes from the fact that Tim9 and Tim10 differ in the sequence length, particularly the number of residues *n* in the CX_3_C-X_*n*_-CX_3_C motif, which differs by one; adding an additional residue in the EM map or removing one results in a less good fit (not shown). Consequently it is likely that the assignment of Tim subunits in the EM maps was correct. Therefore, we speculate that the interactions of the TIM9·10·12 complex within the intact TIM22 complex contribute to stabilizing certain stoichiometries.

We found that the assembled hexameric Tim chaperones are in equilibrium with the unassembled subunits (ca. 10 % population; temperature-dependent, Fig. 3), exchanging on a minutes time scale. Mechanistically, subunit exchange can proceed either via a transient species with less than 6 subunits (i.e., subunits are ejected from the complex before the insertion of new subunits), or via a transient oligomer with more subunits, which then releases subunits. Ivanova *et al*. have studied the kinetic assembly of TIM9·10 by kinetic spectroscopy, and reported intermediates of assembly with four instead of six subunits (25). This species may be involved in the subunit exchange process reported here. The time scale they reported for the tetramer-to-hexamer step at 25 *°*C was of the order of ca. 2 seconds, much faster than the overall exchange rate we observe. However, the reaction we observe comprises disassembly and reassembly reactions, and the net rate of subunit exchange is expected to be lower. No rate constant of disassembly had been reported before, but the equilibrium of populations of free subunits over hexamer found in our study, which is a few percent (Fig. 3C) suggests that the disassembly reaction must be at least one order of magnitude slower than the assembly, and thus i.e. in the tens-of-seconds to minutes range. Our experiments rely on observing incorporation of isotope-labeled subunits into unlabeled hexamer from a mixture, the apparent exchange rate is, thus, reduced by reactions in which the same subunits (unlabeled) re-insert into the hexamer, which are invisible to our approach. Taken together, the minutes time scale we observe is in reasonable agreement with an assembly on a seconds time scale and disassembly on tens-of-seconds to minutes (although somewhat slower). We propose that this subunit exchange is the mechanism by which newly imported Tim subunits get incorporated into pre-existing hexamers, allowing e.g. the formation of TIM9·10·12 from TIM9·10.

For the cell, the continuous presence of a population of unassembled Tim subunits comes at a price: because unassembled Tim subunits are degraded by the IMS AAA+ protease system Yme1 (26), TIM chaperones are continuously removed from the IMS, and the pool of chaperones needs to be continuously replenished. The advantage of this removal/replenishment of small TIMs allows an efficient adaptation of the concentration to the cellular requirement for import systems. Thus, the amount of TIM chaperones can be rapidly adapted by modifying the quantity of newly imported Tim subunits, as the TIM chaperones are continuously degraded through the ca. 10 % of unassembled species.

Interestingly, the functionally related but structurally distinct bacterial periplasmic membrane-protein chaperone Skp also exists in equilibrium between its trimeric functional form and monomers, with a monomer population of ca. 5 % at 37 *°*C (27), suggesting that similar monomer-oligomer equilibria are at work in a variety of chaperones. The ability to rapidly adapt the chaperone concentration likely outweighs the energetic cost related to the need of continuously replenishing the chaperone pool. Furthermore, assembly of chaperones from monomeric subunits may allow the chaperone activation as client proteins emerge (28).

Together with our recent observation that TIM-preprotein complexes are highly dynamic (11), this study stresses the importance of considering molecular dynamics internal motions of molecules as well as dynamic binding/subunit exchange -as a central feature of the mitochondrial TIM chaperone system.

## Methods

### Sample preparation

All proteins used in this study were obtained by recombinant bacterial overexpression. In all cases where isolated small Tims were studied by NMR, the proteins were ^15^N,^13^C labeled, without deuteration, while the labeling pattern for subunits within hexamers included perdeuteration. Isotope labeling was obtained by bacterial growth in M9 minimal medium supplemented with ^15^N-ammonium chloride (1 g/L) and or ^13^C-labeled glucose (2 g/L). For deuterated proteins, ^2^H,^13^C glucose was used, and the growth medium was in 99.9 % deuterium oxide. Unlabeled proteins were obtained by bacterial growth in rich medium (LB), and were used for methods other than NMR, or for NMR samples in which only one subunits was observed.

#### Preparation of Tim9, Tim10 and TIM9·10

Saccharomyces cerevisiae (Sc) Tim9 (MDALNSKEQQEFQKVVEQKQM KDFMRLYSNLVERCFTDCVNDFTTSKLTTNKEQTCI MKCSEKFLKHSERVGQRFQEQNAALGQGLGR) was cloned via NdeI/XhoI into the multiple cloning site (MCS)2 of pETDuet-1, subsequently ScTim10 (MGSSHHHHHHS QDPENLYFQGSFLGFGGGQPQLSSQQKIQAAEAELDL VTDMFNKLVNNCYKKCINTSYSEGELNKNESSCLDR CVAKYFETNVQVGENMQKMGQSFNAAGKF) containing a N-terminal TEV (Tobacco etch virus) cleavage site was cloned via BamHI/HindIII into the MCS1. Tim9 and His-tagged Tim10 were co-expressed in SHuffle T7 *E. coli* cells.

After induction at OD_600_ 0.8 with 0.5 mM isopropyl *β*-D-1-thiogalactopyranoside (IPTG), cells were incubated at 20*°*C overnight and harvested by centrifugation. The cell pellet was resuspended in 25 mL 50 mM Tris-HCl pH 7.4, 150 mM NaCl (buffer A) per liter of cell culture and sonicated. After a heat shock for 15 minutes at 80 ° the soluble fraction was separated from the insoluble fraction by centrifugation at 39121xg for 30 min. The supernatant was loaded on a NiNTA affinity column pre-equilibrated with buffer A, then washed with ten column volume (CV)s buffer A supplemented with 20 mM Imidazole and 0.5 M NaCl (buffer B). Hexameric TIM9·10 complex was eluted with buffer A supplemented with 0.5 M Imidazole (buffer C). The His-tag was cleaved overnight in buffer A supplemented with 0.5 mM dithiothreitol (DTT) and 0.5 mM ethylenediaminetetraacetic acid (EDTA) by adding 1 mg TEV protease to 50 mg protein, cleaved TIM910 was separated from uncleaved TIM910 on a NiNTA column using the above described protocol. The load and wash fractions were then further purified on a HiLoad 16/600 Superdex S200 PG, where TIM910 eluted as a single peak. For NMR experiments gel filtration was performed with 20 mM MES (2-(N-morpholino)ethanesulfonic acid) pH 6.5, 50 mM NaCl (NMR buffer). An alternative method consists of mixing individually expressed and purified Tim9 and Tim10 components, which spontaneously from a complex, as reported (29). Samples produced with any of these methods are indistinguishable in terms of structure and dynamics, as seen by solution-NMR methods (data now shown).

To purify the individual subunits of the TIM9·10 complex, an additional washing step was included into the Ni-NTA purification protocol: after washing with buffer B, the column was washed with ten CVs of buffer B supplemented with 3 M guanidine-HCl to dissociate the TIM910 complex. While His-tagged Tim10 was still bound to the Ni-NTA column, Tim9 was found only in the wash fraction. His-tagged Tim10 was eluted with buffer C supplemented with 3 M guanidine-HCl and refolded by flash dilution into buffer A, the His-tag was cleaved as described above. The wash fraction including Tim9 was dialysed against buffer A and both (Tim9 and Tim10) subjected to gel filtration on a HiLoad 16/600 75 PG. Tim9 was found to elute as a dimer, Tim10 mainly as a monomer. To prepare complex samples in which only one type of subunit was isotope labeled, this procedure was performed twice, starting either from isotope-labeled or unlabeled TIM9·10. The hexamer spontaneously reassembled from Tim9 and Tim10. This property allowed us to prepare complex samples with any desired labeling pattern of the two types of subunits.

All samples of reduced species in this study were prepared by supplementing the NMR buffer with 10 mM tris(2-carboxyethyl)phosphine (TCEP). We have also expressed Tim9 and Tim10 individually, with expression vectors containing only one sequence, but the yield of the pET-Duet expression vector (ca. 60 mg/L) was about one order of magnitude larger than expression of the individual subunits.

#### Preparation of Tim12 and TIM9·10·12

The sequence of Tim12 with an N-terminal His_6_-tag in a pET28a plasmid was purchased from GeneCust (sequence: MGHHHHHHENLY FQGSFFLNSLRGNQEVSQEKLDVAGVQFDAMASTFN NILSTCLEKCIPHEGFGEPDLTKGEQACIDRCVAKMH YSNRLIGGFVQTRGFGPENQLRHYSRFVAKEIADDSK K). In addition to the four cysteines that are highly conserved in small Tims and structurally important (CX_3_C motif), yeast Tim12 contains two additional cysteines, C31 and C61. Initial expression and purification tests with wild-type Tim12 showed a tendency to aggregation. To simplify expression and purification, we replaced these non-conserved cysteines by alanines, resulting in scTim12-C31A-C61A, henceforth called Tim12. Expression of Tim12 was performed in SHuffle T7 *E. coli* cells as described for TIM9·10. Tim12 was found in the insoluble fraction and purified by His-affinity chromatography under denaturing conditions in 6 M guanidine-HCl. Purified Tim12 was bound to a NiNTA column, washed with five CVs of buffer A and incubated with a threefold excess of TIM9·10 for 30 min and subsequently eluted. After TEV-cleavage of the His_6_ tag of Tim12, a gel filtration profile on a HiLoad 16/600 Superdex S200 PG in buffer A shows two separate peaks, Tim12 and Tim12 in complex with TIM910. For the production of TIM9·10·12 samples containing only a single type of isotope-labeled subunit, this procedure was performed either with TIM9·10 composed of unlabeled and labeled subunits, or by using isotope-labeled Tim12 with unlabeled TIM9·10.

### NMR spectroscopy

All NMR experiments were performed on Bruker Avance-III spectrometers operating at 600 MHz ^1^H Larmor frequency (14.1 Tesla). For the resonance assignment of Tim9, Tim10 and Tim12, as well as of Tim12 in TIM9·10·12, the following experiments were performed : 2D 15N-1H-BEST-TROSY HSQC, 3D BEST-TROSY HNCO, 3D BEST-TROSY HNcaCO, 3D BEST-TROSY HNCA, 3D BEST-TROSY HNcoCA, 3D BEST-TROSY HNcocaCB and 3D BEST-TROSY HNcaCB.(30, 31). The sample temperature was set to 25*°*C and sample concentrations were of the order of 200-400 µM. The NMR resonance assignment of TIM9·10 was reported earlier (11). ^15^N R_1_ and R_2_ relaxation rate constants and ^1^H-^1^5N heteronuclear Overhauser enhancements were obtained from series of 2D ^1^H-^1^5N experiments as implemented in NMRlib (32). Twelve relaxation delays were used for T_1_ experiments (0.01 to 1.21 s) and 14 delays (0 to 85 ms) were used for T_2_ measurements. Diffusion-ordered NMR spectroscopy (DOSY) experiments were performed at 600 MHz, from a series of one-dimensional ^15^N-filtered DOSY experiments or (for TIM9·10) a series of two-dimensional ^1^H-^15^N-edited DOSY spectra. Diffusion coefficients were obtained from fitting integrated 1D (excluding the NH_2_ resonances) or 2D spectral intensities as a function of the gradient strength (at constant diffusion delay). The data reported in Fig. 3B was obtained from summing observable minor-state peaks in the 2D series. The kinetics at which subunits are incorporated into hexameric species was followed by observing the buildup rate of NMR peaks corresponding to the hexameric state of a series of ^1^H-^15^N BEST-TROSY HSQC spectra. To probe incorporation of unlabeled TIM910 or TIM910-GGC complex was mixed in a 1:3 ratio with [^2^H,^15^N,^13^C] Tim9 or Tim10. Immediately after mixing (dead time of 3-4 minutes), a series of ^1^H-^15^N-BEST-TROSY HSQC experiments was started for 4 hours whereby one single experiment lasted 3 min 40 s. Data was analysed by following the build up of TIM910 complex peaks or the loss of signal intensity in Tim9/Tim10 free subunit peaks. The sum of the peak intensity at each time point was plotted against the time and a mono exponential fit performed.

### Analytical ultracentrifugation

Sedimentation velocity experiments were performed on the Biophysical Platform AUC-PAOL at IBS Grenoble, on an analytical ultracentrifuge XLI, with a rotor Anti-50 (Beckman Coulter, Palo Alto, USA) and double-sector cells of 12 mm and 1.5 mm optical path length equipped with Sapphire windows (Nanolytics, Potsdam, Germany). Samples were measured in NMR buffer at 10*°*C at three different concentrations from 0.2 to 0.002 mM, and interference and absorption at 277 nm wavelength were recorded. The data were processed by Redate software v 0.2.1. The analysis was done with the SEDFIT software, version 14.7 g and Gussi 1.1.0a6.

### Small-angle X-ray scattering

SAXS data collection followed the methods we reported earlier(11). Briefly, data were collected at the beam line BM29 at ESRF(33) with a Pilatus 1M detector (Dectris) at the distance of 2.872 m from the flow-through capillary, using the scattering of pure water for calibrating the intensity to absolute units. The X-ray energy was 12.5 keV and the accessible q-range 0.032 nm^*−*1^ to 4.9 nm^*−*1^. The incoming flux at the sample position was in the order of 1012 photons/s in 700×700 *µ*m^2^. Automatical azimuthal averaging of the images was done with pyFAI.(34) TIM9·10·12 was subjected to a gel filtration directly coupled to the SAXS beamline (see Figure S3D). Exposures with radiation damage were discarded, the remaining frames averaged and the background was subtracted by an online processing pipeline.(35) Data from the three concentrations were merged following standard procedures to create an idealized scattering curve, using Primus from the ATSAS package.(36) The pair distribution function p(r) was calculated using GNOM.(37) Example data of the SEC-SAXS experimentsare shown in Fig. S3D.

### Molecular Dynamics simulations

First we used the structure of the TIM9·10 hexamer built in our previous work (11) as the initial model of the Tim9_2_·Tim10_3_·Tim12 hexamer. Then one of the Tim9 subunits was replaced by a Tim12 subunit which was built using the crystal structure of TIM9·10 (PDB 3DXR (12)) as the templates based on the sequence of yeast Tim12 (UniProtKB: P32830). All missing residues were modelled using MODELLER9.18 (38). The disulfide bonds present in the crystal structures were kept.

The TIM9·10·12 hexamer was then placed into a periodic cubic box with sides of 14.2 nm solvated with TIP3P water molecules containing *Na*^+^ and *Cl*^*−*^ ions at 0.10 M, resulting in ∼378,000 atoms in total. The Amber ff99SB-disp force field (39) was used for the simulations. The temperature and pressure were kept constant at 300 K using the v-rescale thermostat and at 1.0 bar using the Parrinello-Rahman barostat with a 2 ps time coupling constant, respectively. Neighbor searching was performed every 10 steps. The PME algorithm was used for electrostatic interactions. A single cut-off of 1.0 nm was used for both the PME algorithm and Van der Waals interactions. A reciprocal grid of 96 x 96 x 96 cells was used with 4th order B-spline interpolation. The hydrogen mass repartitioning technique (40) was employed with a single LINCS iteration (expansion order 6) (41), allowing simulations to be performed with an integration time step of 4 fs. MD simulations were performed using Gromacs 5.1 or 2018.1 (42). The all-atom simulation revealed the *α*-helices of the TIM9·10·12 hexamer are relatively rigid and the tails are flexible towards the N- and C-terminus.

A total of ∼3 *µs* trajectories were collected to sample the conformational space of the TIM9·10·12 hexamer. These sampled conformations were used for further ensemble refinement using the Bayasian Maximum Entropy (BME) method (**?**) guided by experimental SAXS data as decribed in our previous work (11). The workflow of the SAXS guided ensemble refinement is briefly described as following: (1) First, we collected the conformational pools of the TIM9·10·12 hexamer by the atomistic MD simulations. (2) We back-calculated the SAXS curve of each frame using Crysol3 (43) and compared the averaged SAXS data with experimental measurement. (3) We employed the BME approach (44, 45) to refine the simulated ensembles so as to improve the fitting with experimental SAXS data.

## Supporting information

Movie showing an MD simulation of TIM91012

## Acknowledgements

We thank Aline Le Roy and Christine Ebel for excellent support with AUC experiments, and Nils Wiedemann, Caroline Lindau, Beate Bersch and Vilius Kurauskas for insightful discussions, and Bernhard Brutscher, Alicia Vallet and Adrien Favier for excellent NMR platform operation. This study was supported by the European Research Council (StG-2012-311318-ProtDyn2Function), the Agence Nationale de la Recherche (ANR-18-CE92-0032), and the Lundbeck Foundation BRAINSTRUC initiative. This study used the platforms (NMR, EM, isotope-labeling) of the Grenoble Instruct-ERIC Center (ISBG; UMS 3518 CNRS-CEA-UJF-EMBL) with support from FRISBI (ANR-10-INSB-05-02) and GRAL (ANR-10-LABX-49-01) within the Grenoble Partnership for Structural Biology (PSB). The experiments were performed on beamline BM29 at the European Synchrotron Radiation Facility (ESRF), Grenoble, France. The simulations were performed on the Danish National Supercomputer for Life Sciences (Computerome).

## Author contributions

K. W. and P. S. designed and initiated the research project. K.. and A. H. prepared all in-vitro protein samples. K. W. and P. S. performed and analyzed NMR experiments. K. W. and M. B. performed and analyzed the SAXS data. Y. W. and K. L.-L. designed and performed the MD simulations. K. W., Y. W.and P. S. prepared the figures. P. S. wrote the paper with input from all authors.

## Data availability

The chemical shift assignments of Tim9, Tim10 and Tim12 have been tabulated as TALOS input files and are available on Mendeley data: http://dx.doi.org/10.17632/b3n93f3bcr.2. The MD model and the SAXS data have been deposited in the SASBDB (accession number SASDH28) and is available at https://www.sasbdb.org/data/SASDH28/mfyw9wkrhr/.

---

**Fig. S1.**
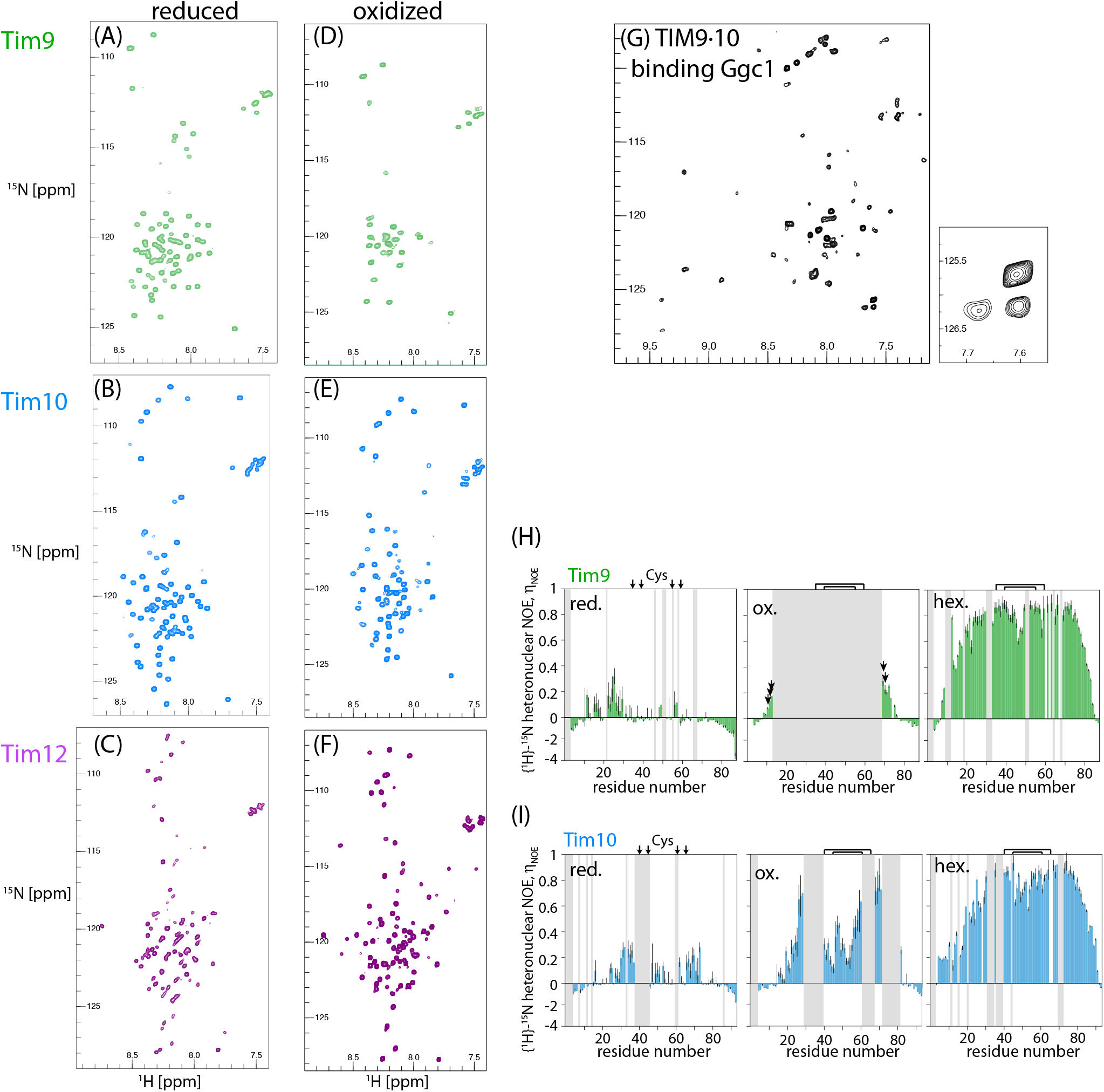
^1^H-^15^N correlation spectra of all samples described in this study, in the isolated Cys-oxidized state (i.e. without reducing agent; (A-C)) or Cys-reduced state (with tris(2-carboxyethyl)phosphine; (D-F)). (H) TIM9·10 bound to the GDP/GTP carrier Ggc1. As described elsewhere (11), the increased size (two hexamers bind one Ggc1 chain) and presence of millisecond dynamics in the binding cleft lead to strong intensity reduction. (H,I) ^1^H-^15^N heteronuclear Nuclear Overhauser effects in the various monomeric and hexameric Tim9 and Tim10 states. Error bars were estimated from three times the spectral noise level.

**Fig. S2.**
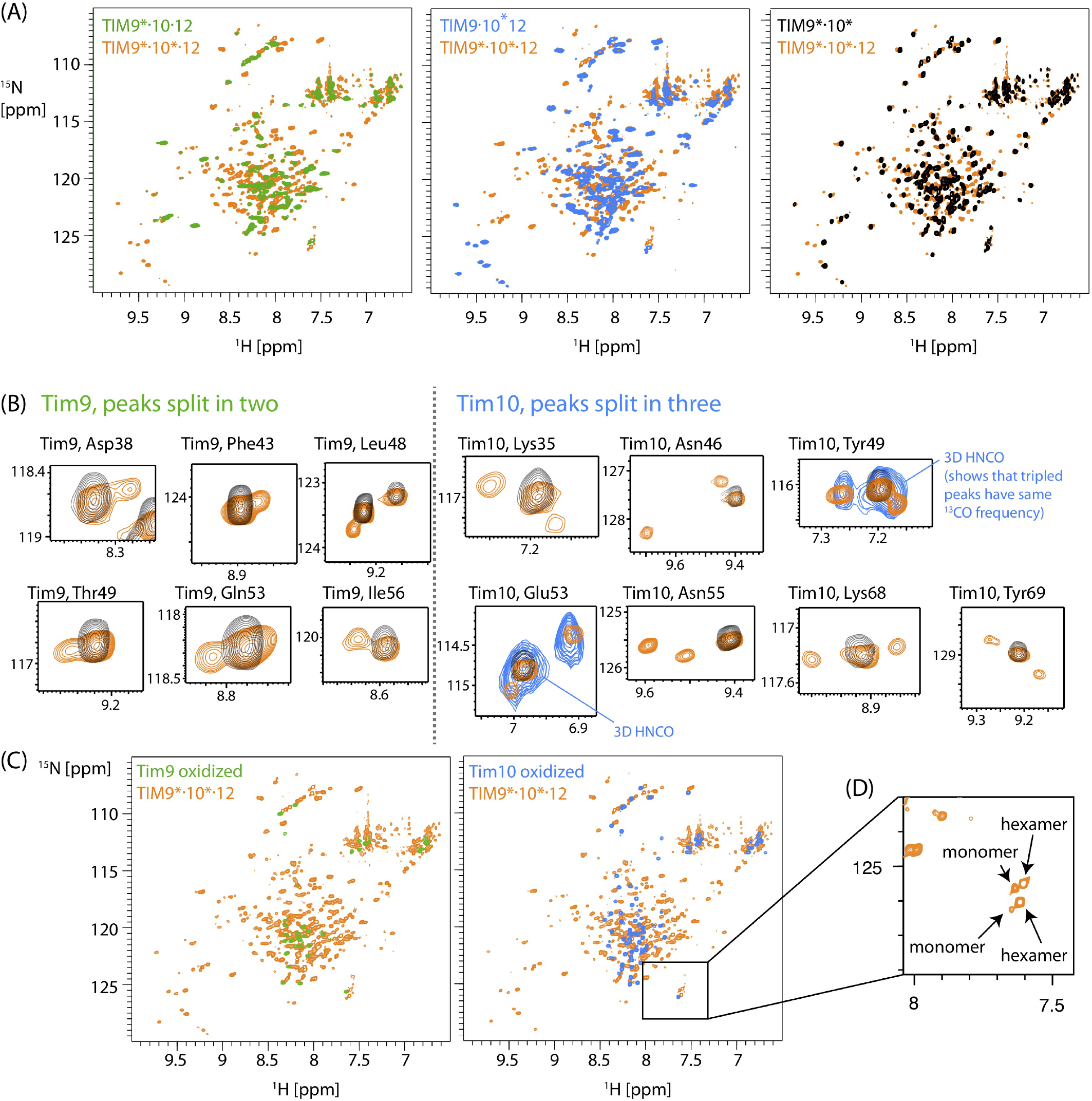
NMR spectra (^1^H-^15^N BEST-TROSY spectra) of the TIM9·10·12 complex with different isotope-labeling schemes, and comparison to spectra of TIM9·10 as well as the disassembled subunits. (A) The spectrum in which Tim9 and Tim10 subunits are labeled, within the TIM9·10·12 complex, is shown in orange and overlaid with a spectrum in which only one subunit is labeled (green: Tim9; blue: Tim10). The spectrum of the TIM9·10 complex, in which both subunit types are labeled, is shown in black (right). (B) Zoom into the 2D spectra of TIM9·10·12 and TIM9·10, exemplifying the observed peak multiplicity observed for residues of Tim9 (left) and Tim10 subunits (right). In order to confirm that the tripled peaks indeed stem from the same residue in three different environments, we collected a 3D HNCO spectrum with the TIM9·10·12 sample in which only Tim10 is isotope-labeled. This spectrum is overlaid, at the corresponding carbonyl frequency, on the 2D spectra (for readability only for two examples) and confirms the assignment of the three peaks to the same residue. Evidence for peak duplication is observed for many residues of Tim9, but no peak triplication is observed. Peak triplication is observed exclusively for residues belonging to Tim10. (C) Overlay of the TIM9·10·12 spectrum (only Tim9 and Tim10 labeled) and the spectra of the isolated subunits (Tim9: green; Tim10: blue). The peaks of the isolated subunits do not coincide with the additional peaks observed in TIM9·10·12 and not in TIM9·10. This observation demonstrates that the additional peaks in TIM9·10·12 relative to TIM9·10 do not stem from isolated subunits in equilibrium with hexamers, but must correspond to distinct conformations within the TIM9·10·12 hexamer. (D) Zoom onto the cross-peaks of the two C-terminal residues in Tim9 and Tim10, respectively, showing that also in TIM9·10·12 there is a minor form, corresponding to free subunits.

**Fig. S3.**
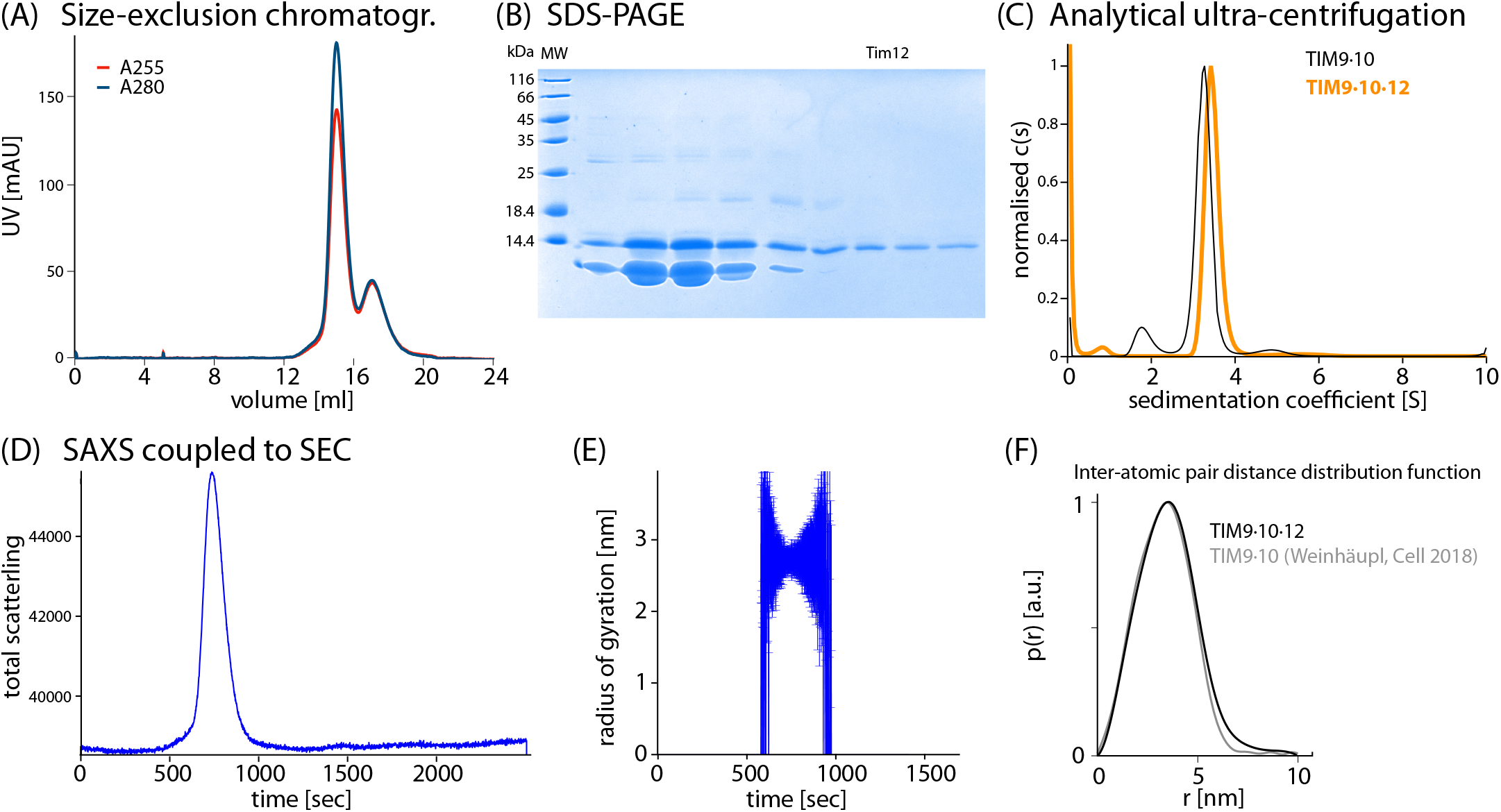
Biochemical and biophysical characterization of TIM9·10·12. (A) Gel filtration profile of the sample following the complex formation procedure described in the Methods section. The main peak contains Tim9, Tim10 and Tim12; re-injection of the fraction of this peak into a size-exclusion chromatography results in only a single peak, i.e. the TIM9·10·12 hexamer remains stable (see also panel (E)). (B) Gel electrophoresis of the fractions from the gel filtration. (C) Analytical ultra-centrifugation profiles of TIM9·10·12, compared to the one of TIM9·10.(11) (D, E) Data from the inline size-exclusion chromatography/small-angle X-ray scattering experiment. The purified TIM9·10·12 complex (main peak in (A) was injected into SEC-SAXS(46) and the scattering was recorded (D) over time during the gel filtration. Panel (E) shows the radius of gyration from the analysis of the SAXS data along the gel filtration, showing that the sample is homogeneous across the entire peak. Note the large error bars towards the extremities of the SEC peak, where sample amount and thus detection sensitivity is low. (F) Comparison of pair distance distributions of TIM9·10·12 with previously reported data of TIM9·10 (11).

**Fig. S4.**
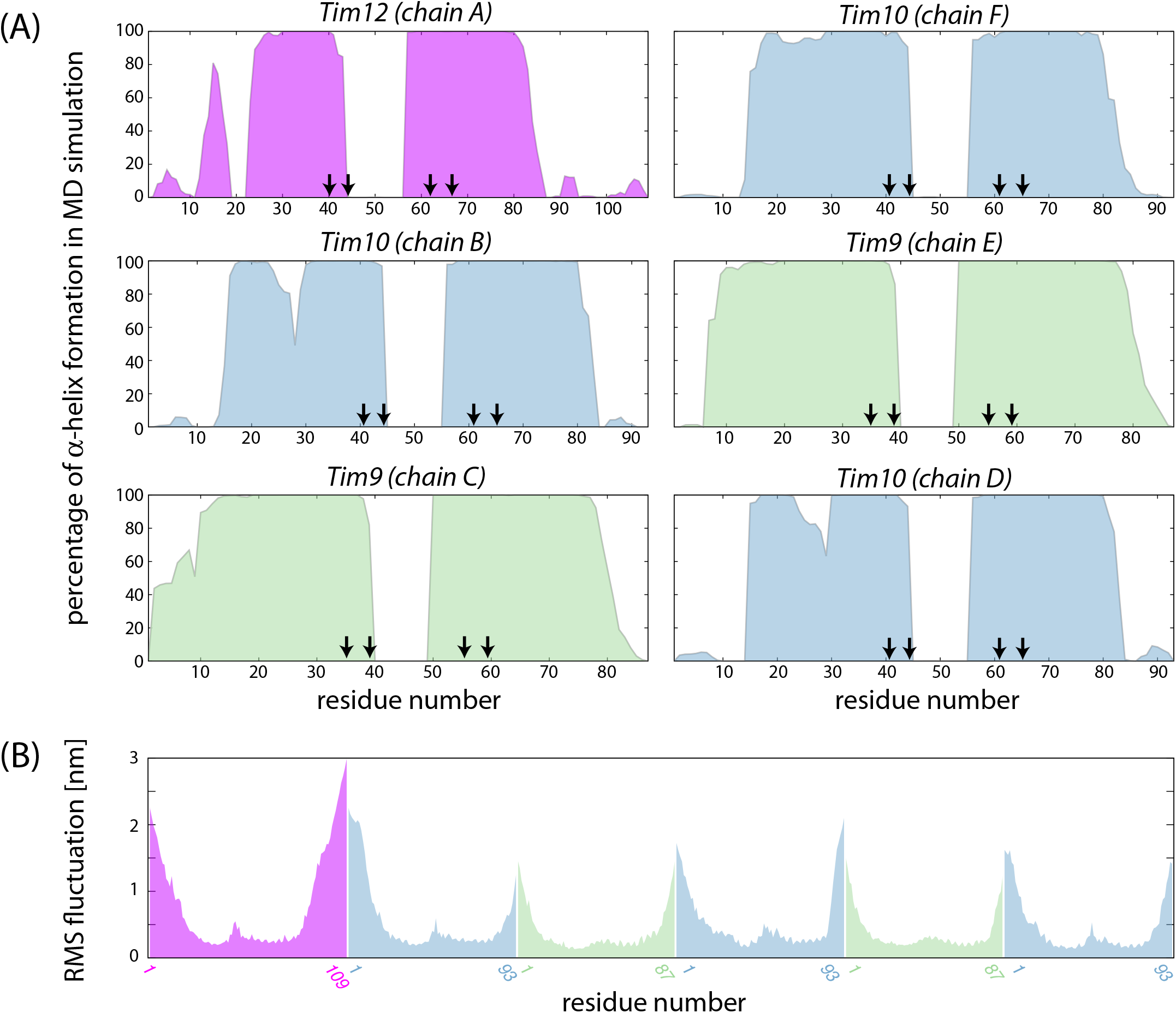
Propensity for formation of helical structure (A) and room-mean-square fluctuation (B) in TIM9·10·12 obtained from the molecular-dynamics simulation. In (A), vertical arrows denote the positions of the disulfide-forming cysteines.

**Fig. S5.**
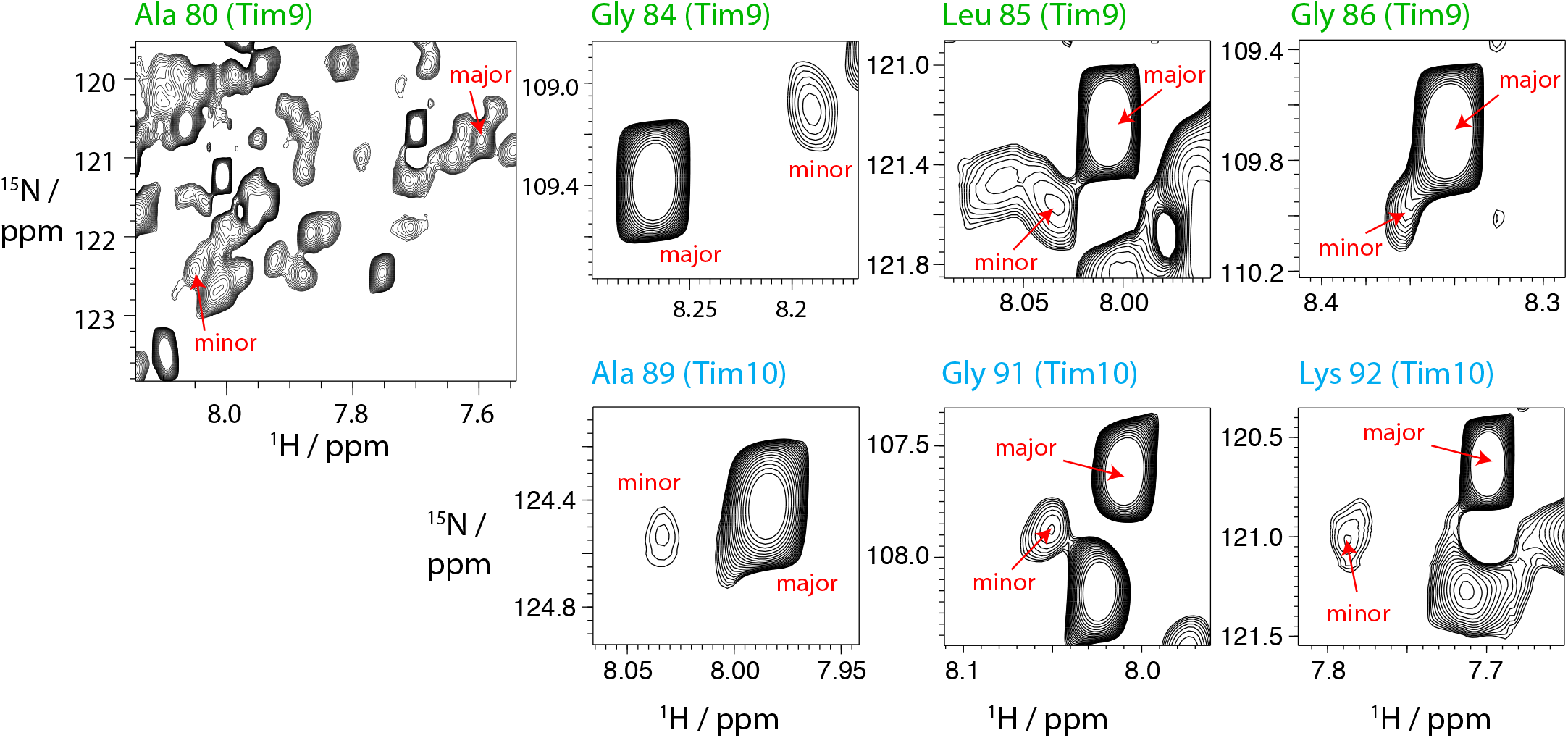
Zoom onto ^1^H-^15^N correlation peaks of selected residues in TIM9·10, highlighting the minor peaks, that correspond to free subunits in equilibrium with the hexamer peaks.

